# Optimising crown-of-thorns starfish control effort on the Great Barrier Reef

**DOI:** 10.1101/2024.04.10.588969

**Authors:** Kanupriya Agarwal, Michael Bode, Kate J. Helmstedt

## Abstract

Outbreaks of crown-of-thorns starfish *Acanthaster planci* (COTS), a disruptive coral-eating predator, are responsible for almost half of total coral cover loss on Australia’s Great Barrier Reef. As the pressures of climate change continue to intensify the frequency and severity of disturbance events such as cyclones and coral bleaching, efficiently managing COTS outbreaks is essential for reef protection. We aim to understand how the spatial distribution and intensity of crown-of-thorns starfish control – specifically manual culling of COTS by human divers – can impact coral cover on the GBR. We construct a metapopulation model based on a predator-prey model with larval dispersal and removal of crown-of-thorns starfish to simulate and compare spatial control strategies. When outbreaks begin on reefs between Cairns and Cooktown, we found the best strategy is to target those reefs at the source of the COTS outbreak. Increasing the spatial spread of control results in a larger spatial area protected across the GBR, but a lower total coral cover on the GBR. Our findings suggest that carefully targeting future control by considering larval connectivity patterns and spatial control strategies could lead to more efficient crown-of-thorns management. With the increasing pressures of climate change, any efficiency gains in reef management will prove beneficial for the Great Barrier Reef.

## 1 Introduction

Coral reefs such as Australia’s Great Barrier Reef (GBR) are among the most diverse ecosystems on Earth, supporting approximately 25% of all marine species in approximately 0.1% of its area (Knowlton et al., 2010; Fisher et al., 2015). Yet, in the 27-year period between 1985 and 2012, the GBR’s coral cover declined by over 50% (De’ath et al., 2012; Dietzel et al., 2020). Outbreaks of crown-of-thorns starfish *Acanthaster planci* (COTS), a disruptive coral-eating predator, are a major driver of coral reef degradation, responsible for 42% of this loss of coral cover on the GBR (De’ath et al., 2012; Osborne et al., 2011). These corallivorous starfish compound the negative impacts of climate change, including cyclones and coral bleaching (Hughes et al., 2003, 2017). Therefore, efficiently managing COTS outbreaks is essential for reef protection, as the pressures of climate change continue to increase the frequency and severity of disturbance events (Osborne et al., 2017).

Managing crown-of-thorns starfish outbreaks is difficult due to the local density of COTS populations on outbreak reefs, and their potential presence across hundreds of reefs and tens of thousands of square kilometres. Indirect actions including marine reserve zoning and improving water quality can impact COTS populations, but the most direct control action is manual culling of individual COTS with lethal injections by human divers (Westcott and Fletcher, 2018). At small local scales, manual control of COTS has varying levels of success (Westcott and Fletcher, 2018), since even identifying and detecting COTS can be challenging (Heenaye-Mamode Khan et al 2023). However, coordinated large-scale control programs can effectively reducing COTS densities (Fletcher et al., 2020; Matthews et al., 2024). The GBR consists of nearly two thousand connected reefs, so a thorough analysis of COTS control must be considered on a whole-of-GBR scale (Hock et al., 2014; Vanhatalo et al., 2017). A range of statistical and simulation models have been used to model crown-of-thorns starfish dynamics across the GBR, but comparative analyses of control strategies are rare (Castro-Sanguino et al. 2023; Hock et al. 2014; Fletcher et al. 2021). To undertake a comprehensive analysis of COTS control, we need (1) dynamic large-scale models that can incorporate larval dispersal (integral to coral and COTS population dynamics), (2) models of how COTS control efforts impact densities, and (3) optimisation methods that can compare a vast number of potential spatial control strategies.

Models of COTS control across the Great Barrier Reef focus on faithfully representing the spatial ecology of outbreaks. For example, the connectivity and larval recruitment of COTS and coral, or on the predator-prey dynamics of COTS outbreaks on individual coral reefs (Condie et al., 2012; Hock et al., 2017; Vanhatalo et al., 2017). These models assume that control efforts will follow current “priority reef” strategies (Fletcher et al. 2021; Castro-Sanguino et al. 2023). These investigate the effects of current COTS control efforts, however they focus their control in a fixed same spatial area rather considering than the need to spread or focus a fixed control budget over different control area sizes.

We investigate the impact of various crown-of-thorns starfish control strategies on COTS and coral populations on the Great Barrier Reef. We develop a metapopulation model for the GBR, based on Morello et al. (2014), which incorporates larval dispersal for both coral and crown-of-thorns starfish, as well as manual culling of crown-of-thorns starfish. We then simulate a crown-of-thorns starfish outbreak for 1705 reefs on the GBR using this metapopulation model. Finally, we simulate various COTS control strategies during an outbreak. We vary the control strategies by varying the spatial spread and intensity of manual culling across the GBR to answer the question: should control intensity be spread or focused?

## 2 Methods

We develop a model which describes the predator-prey interactions between crown-of-thorns starfish (predator) and coral (prey), and incorporates larval dispersal of crown-of-thorns starfish, larval dispersal of coral, and culling of individual crown-of-thorns starfish by human divers at a large number of distinct reefs. The model accounts for area and position of reefs, and connectivity between them. We use an existing and well-known predator-prey model for a single population from Morello et al. (2014) and extend this model to a predator-prey metapopulation model with larval dispersal and crown-of-thorns starfish control. Finally, we parameterise this model for the Great Barrier Reef and various crown-of-thorns starfish control scenarios.

### 2.1 Predator-prey metapopulation model

The model of intermediate complexity from Morello et al. (2014) includes a discrete age-structured population model for COTS, a logistic growth model for fast-growing coral, and a logistic growth model for slow-growing coral. We only include fast-growing coral in our model, since these are the preferred food source of COTS; we omit slow-growing coral abundance which has less effect on COTS growth and survival (Birkeland and Lucas 1990). The age-structured model for COTS has 3 age classes: age 0 (larvae), age 1 (juveniles), and age 2 or older (adults). We extend the discrete time, single reef model in Morello et al. (2014) to a metapopulation model using dispersal described in Hock (2017), which simulates dispersal between all relevant reefs in the GBR. The predator-prey metapopulation model equations for coral and COTS are as follows:

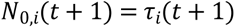

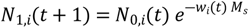

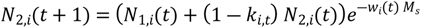

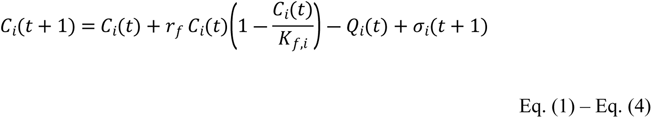

where,

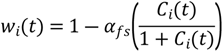

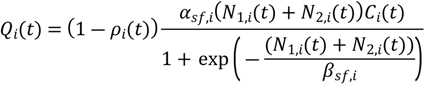

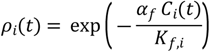

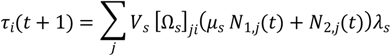

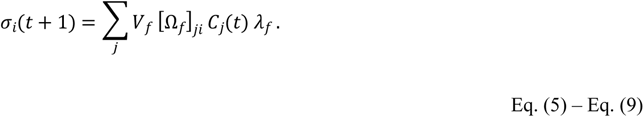

Here *N*_*a,i*_(*t*) denotes the number of COTS of age *a* = 0, 1, 2, at reef *i* in year *t* (note *N*_2,*i*_(*t*) combines all COTS aged over 2). Equations 1 – 3 describe the age-structured population model for COTS with population growth (a function of coral abundance, Equation 5), demographic mortality, and mortality from manual culling. Equation 1, describing COTS of age 0, relies upon connectivity with other reefs in the metapopulation for reproduction via dispersal. Fast-growing coral cover at reef *i* in year *t* + 1, *C*_*i*_ (*t* + 1), is modelled in square kilometres (Equation 4) and incorporates demographic growth and mortality, consumption by COTS (via Equation 6), and reproduction via dispersal. Larval recruitment for both COTS (Equation 8) and coral (Equation 9) are driven by connectivity matrices Ω_*s*_ and Ω_*f*_ respectively, the per-capita fecunditiy of each, and survival rates after settlement. See Supplementary Methods for full description of the model.

### 2.3 COTS culling

We model the manual removal of adult COTS as a percentage of the total population at a given reef. Parameter *k*_*i,t*_ denotes the proportion of age 2+ COTS that are culled at reef *i* in year *t*, and these values *k*_*i,t*_ will be chosen in our simulations. Here, we are modelling population-dependent culling and assuming that control effort has a linear relationship with the proportion of COTS culled.

### 2.4 Parameterising the GBR-wide model

Our GBR model contains 1705 reefs. Where possible, we use parameter values that were calibrated by Morello et al. (2014) for historical coral and COTS populations on the GBR (see Table S1). The connectivity matrices for both COTS and coral larval dispersal were simulated by a high-resolution biophysical model of larval connectivity for GBR reefs between 1996-2002 (Bode et al. 2012). In 1995, a large outbreak began on the reefs around Lizard Island in the northern GBR; by 2002, the outbreak had moved more than 500 km south, to the reefs between Townsville and Mackay in the central GBR. We use the same connectivity matrices for coral and COTS, a common assumption (Hock et al., 2017) that overlooks some behavioural and reproductive differences between the organisms. The GBR reef sizes are sourced from the Great Barrier Reef Marine Park Authority (1998).

We estimated the proportion of age 1 COTS that can reproduce using a model from Lucas (1984) which describes the relationship between COTS age and size, and a model from Babcock et al. (2016) which describes the relationship between COTS size and gonad weight, which we assume scales linearly with reproductive output (see Supplementary Methods for more details). Survival rates in the larval and settlement life stages are very difficult to estimate for both COTS and coral. We therefore combined these processes into a single survival parameter: calibrated to ensure that coral cover did not approach exponential increase in the absence of COTS, and the COTS populations did not approach exponential increase, but still at a relatively high rate in the presence of coral.

We initialise coral cover at 50% on each reef. Although this is higher than the current state of the GBR (where average coral cover ranges between 10-50%, and varies regionally (Australian Institute of Marine Sciences, 2021)), by simulating a COTS outbreak with abundant coral cover across the GBR, we are better able to discriminate between different control actions.

To simulate a COTS outbreak on the GBR, for initial adult COTS populations, we initially place 100 COTS at every reef in the outbreak initiation box (the region between Cairns and Cooktown where COTS outbreaks begin on the GBR; see Supplementary Methods) and model the whole GBR for 100 years. To calculate the initial age 1 (*N*_1,*i*_(0)) and age 0 COTS (*N*_0,*i*_(0)), we recreate a stable age distribution following Morello et al. (2014):

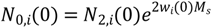

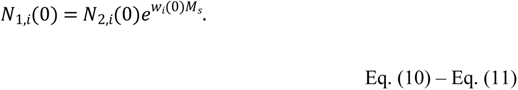

### 2.5 Control scenarios

We simulated four different COTS control scenarios, each distributing a fixed control budget across a differently-sized area in the GBR. The simulation was run with and without control for 100 years, to ensure that both short-term and decadal scale consequences become apparent (Condie et al., 2018). The four scenarios varied the distribution and intensity of control actions, which remained consistent throughout each simulation (Figure 1). We define the total annual control effort for each scenario as the sum of the control effort at every reef,

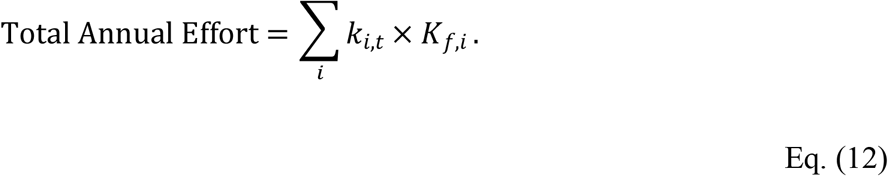

**Figure 1:**
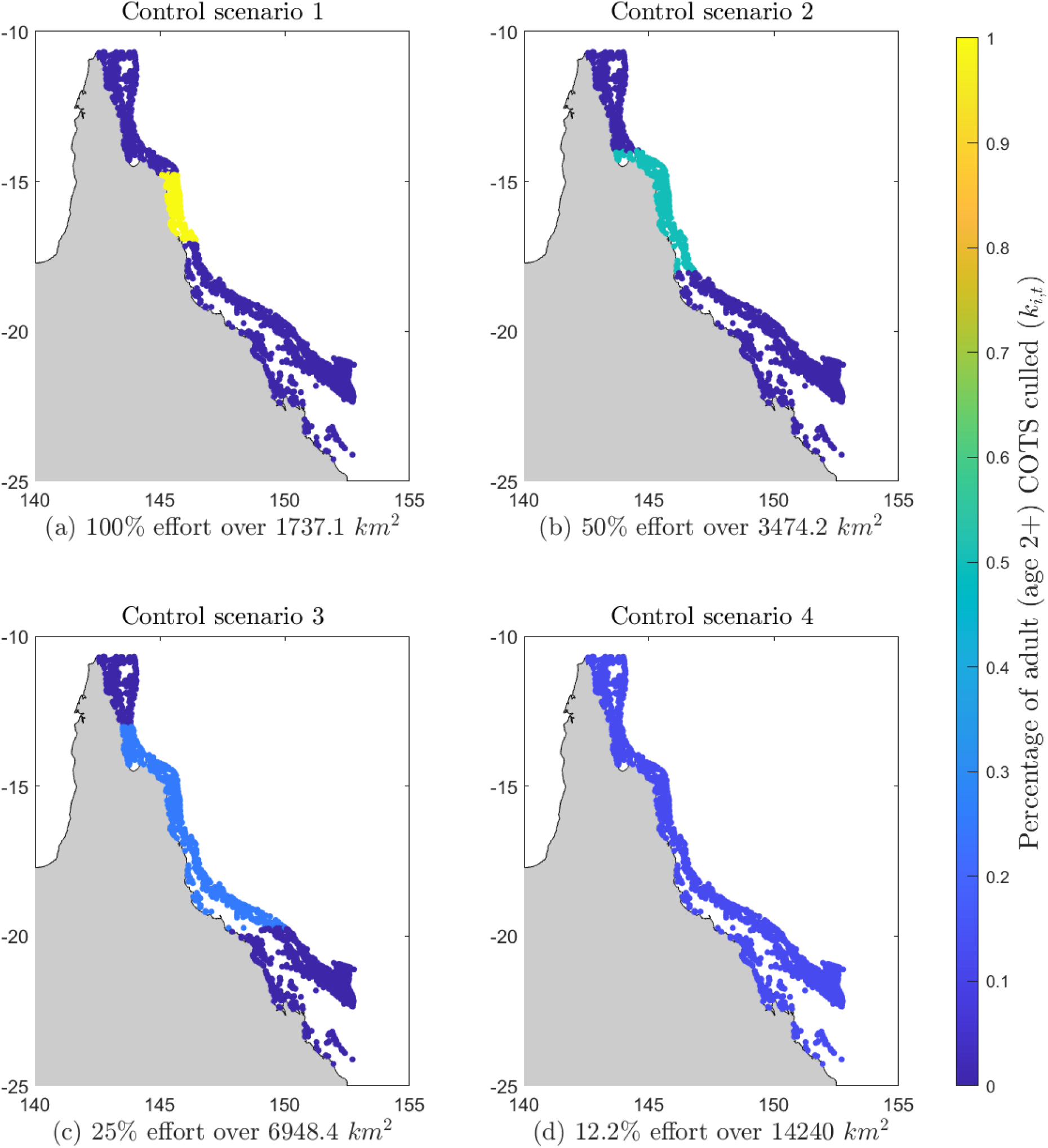
The spatial spread and intensity of COTS control effort across the GBR reefs, for the four scenarios. Each of 1705 individual reefs is indicated by a dot in the map, indicating location but not size. Control effort is quantified by the percentage of adult (age 2+) COTS culled at reef i each year for simulations of 100 years, indicated by colour of the reef (yellow indicating high effort, dark blue indicating low or no effort)..

Each control scenario has the same total annual effort, distributed differently through space. More vessels used for COTS control result in better outcomes (Fletcher et al. 2021), and therefore spreading a fixed budget across a larger area will erode the impact of control at any individual reef. Control science and strategies are rapidly evolving for COTS in the GBR and beyond (Babcock et al. 2020), and so we do not model the control method in detail (see Fletcher et al. 2021, Castro-Sanguino et al. 2023 for assessment of current control strategies), instead assuming a percentage of the adult COTS can be culled across the area. In scenario 1, we cull 100% of adult COTS, which is equal to *k*_*i,t*_ = 1, at every reef in the initiation box, a total reef area of 1737 *km*^2^ (Figure 1a). This scenario attempts to halt the outbreak before it spreads beyond the initiation box (Babcock et al. 2020). In scenario 2, we halve the percentage of COTS culled but double the area controlled, so we cull 50% of adult COTS, or *k*_*i,t*_ = 0.5, over a total reef area of 3474.2 *km*^2^ (Figure 1b). In scenarios 3 and 4, we further increase the number of reefs being treated, each with lower levels of control. By the final scenario, all reefs in the metapopulation are being treated with some control effort. Table 1 describes the details of all four control scenarios including the percentage of adult COTS culled at each reef, the number of reefs controlled, the total area controlled, and the figure which shows the spatial spread of the control effort. We chose the reefs controlled in all scenarios to be centred around the outbreak initiation box since that is the source of the outbreak (Figure 1).

**Table 1:**
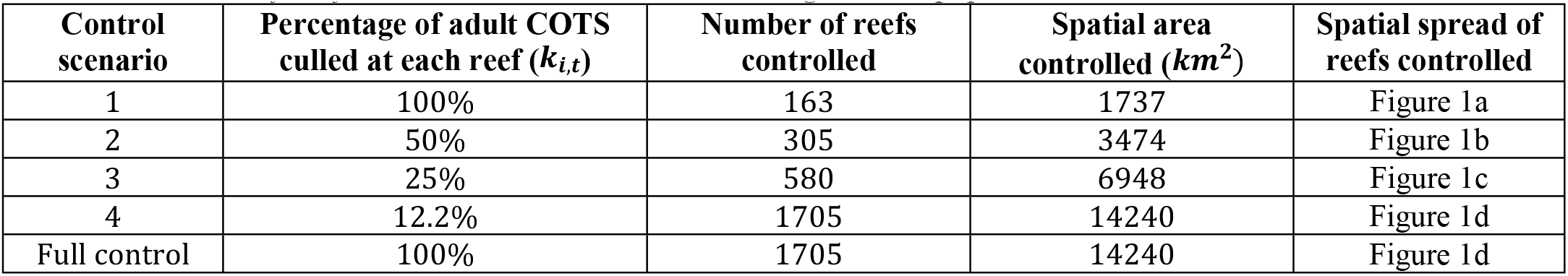
The details of the four control scenarios simulated using our metapopulation model.

We included an additional control scenario with 100% of COTS culled over the entire GBR. This scenario does not have the same total annual effort as the other strategies described in Table 1, and its effectiveness was not directly compared to the other control scenarios. This scenario is not intended to reflect a realistic control strategy, but is used to establish an upper limit for coral cover increases.

To evaluate the effectiveness of each control strategy, we compare the total area of coral cover across all reefs. This measure assumes that management goals are to maximise average coral cover across the GBR. However, other goals include minimising the number of reefs with low coral cover (e.g., a “minimax” strategy), or maximising the number of reefs with the highest coral cover (“maximin”; Peterson, 2017).

## 3 Results

### 3.1 Crown-of-thorns starfish outbreak dynamics without control

Without control, 100 years from the start of the outbreak the crown-of-thorns starfish population spreads to 1700 of the 1705 reefs on the GBR (Figure 2a). However, only 177 reefs achieve high adult COTS populations of 500 or more (Figure 3b). After 100 years, most COTS are not in the outbreak initiation box - only 10 reefs within the outbreak initiation box have 500 or more adult COTS. Instead, the outbreak spreads both north and south on the GBR, and there are two clusters (one north and one south) which have high adult COTS populations (Figure 2). In the north, there is a cluster of 94 reefs with 500 or more adult COTS each, and in the south there is a cluster of 73 reefs with 500 or more adult COTS each. The northern cluster has higher populations of adult COTS – one with 13,649 adult COTS (Figure 2a, darkest purple).

**Figure 2:**
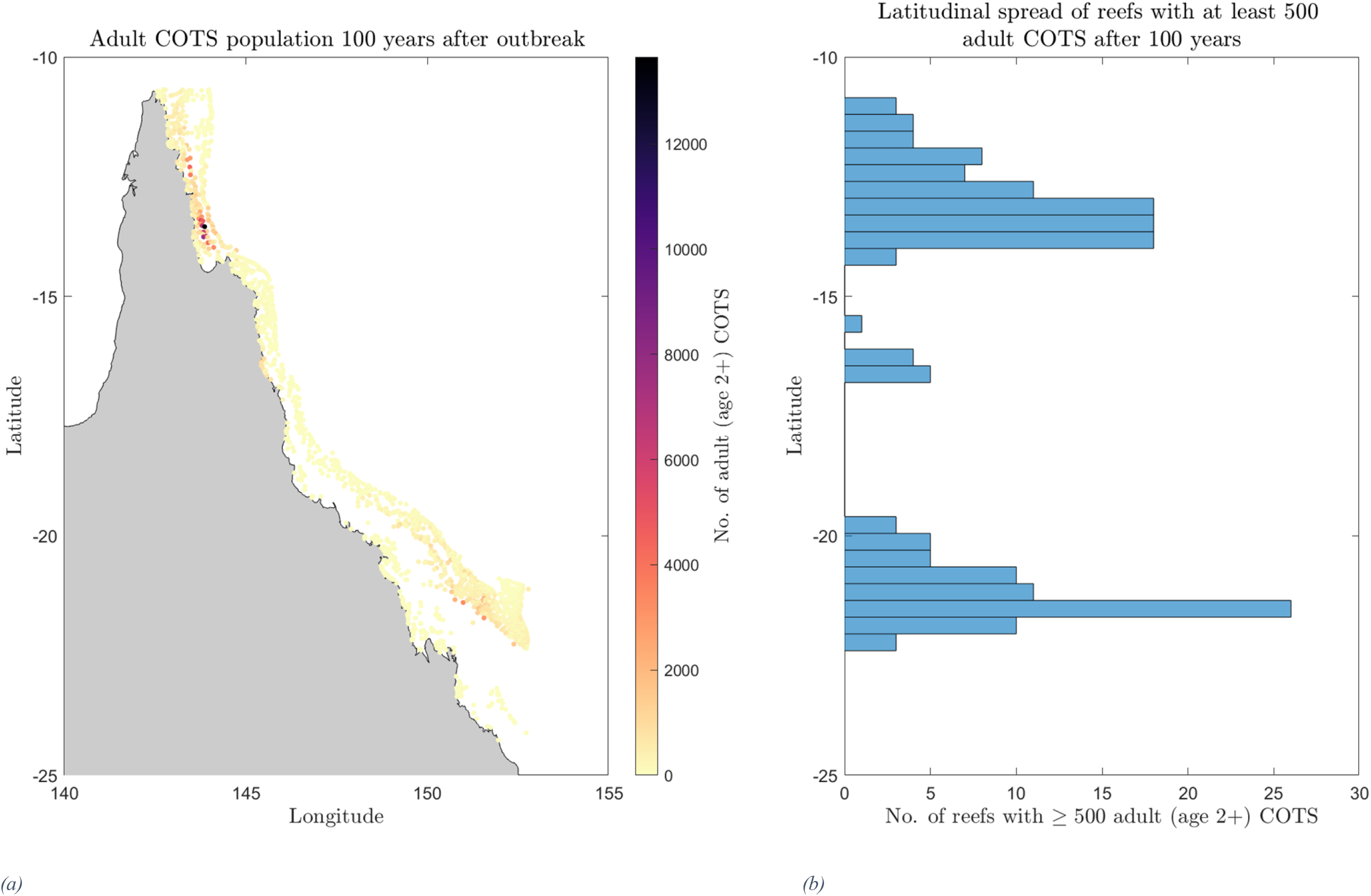
The spatial spread and size of the adult (age 2+) COTS population across the Great Barrier Reef (GBR) after 100 years with no COTS control effort. (a) Map of the GBR, where the colour of each reef corresponds to the number of adult COTS. (b) Latitudinal spread (north-south spread) and count of the reefs on the GBR with at least 500 adult COTS.

**Figure 3:**
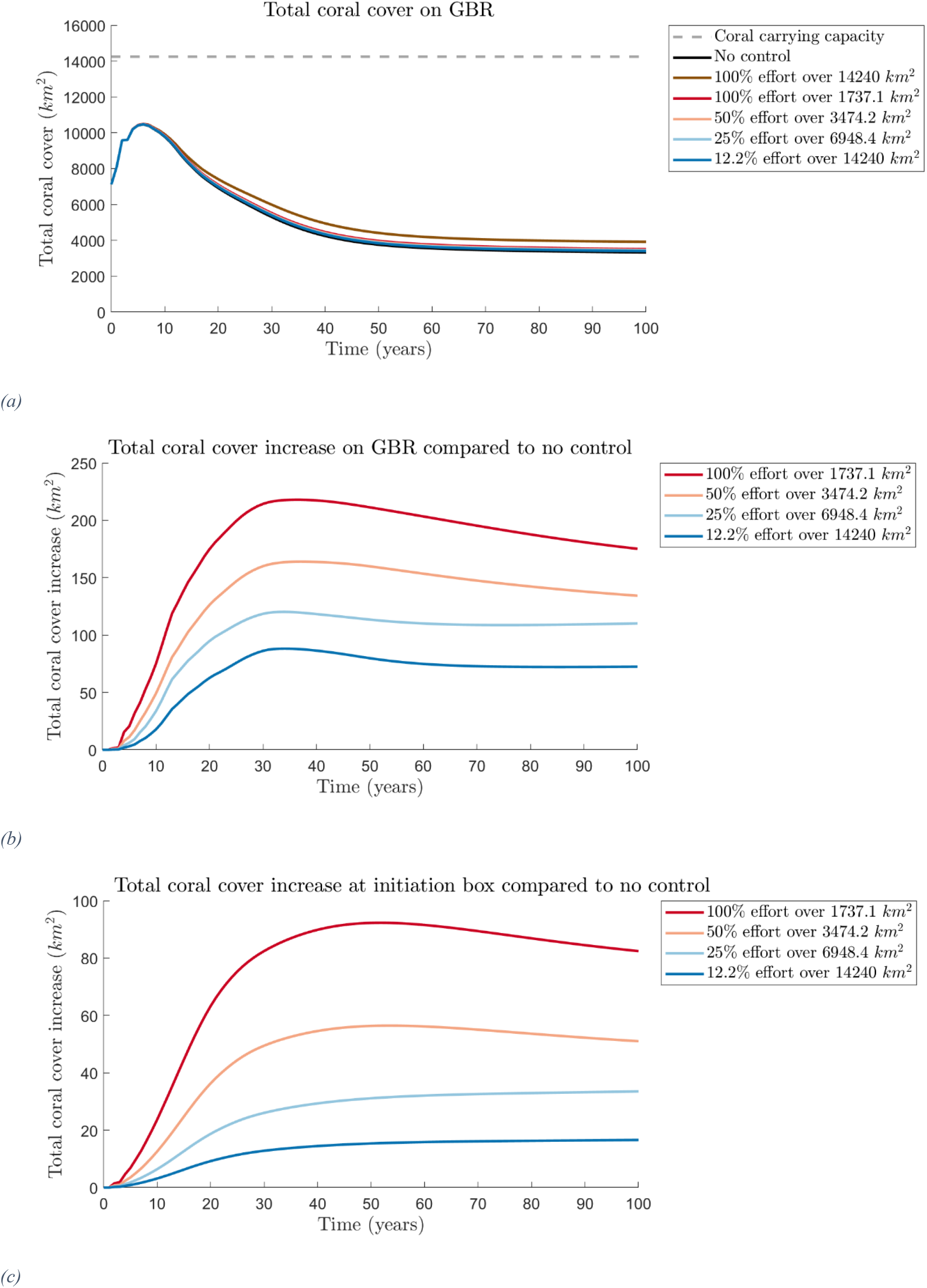
Impacts of COTS control strategies on coral over 100 years. (a) Total coral cover on the GBR, (b) total coral cover increase on the GBR compared to no control effort and (c) total coral cover increase at the initiation box compared to no control effort (in km^2^). Six strategies are shown: 100% control effort over reefs in the initiation box (red), 50% control effort over 3474.2 km^2^ (light red), 25% control effort over 6948.4 km^2^ (light blue), and 12.2% control effort over all reefs (dark blue), no control (black, (a) only), 100% control at all reefs (brown, (a) only).

### 3.2 Comparison of crown-of-thorns starfish control scenarios

The total coral cover on the GBR follows the same general trend over 100 years for all scenarios simulated, including for no control effort (Figure 3a). Coral cover increases initially, while the COTS population is still relatively small and localised, decreases for approximately 50 years as the outbreak spreads through the GBR, and then stabilises for the last 40-50 years, with COTS becoming endemic. A net reduction of coral cover after 100 years is observed in all control scenarios – even culling 100% of COTS across all reefs (Figure 3a). However in comparison to a scenario with no control, the four control scenarios with the same total annual effort achieve modest increases in total coral cover at the end of the 100 years (between 72 km^2^ and 175.2 km^2^ additional live coral cover) (Figure 3b). We see that culling a higher percentage of COTS over a smaller area is more effective at increasing the total coral cover on the GBR, compared to no control, than culling a smaller percentage of COTS over a larger area.

The most effective control strategy is to cull 100% of adult COTS over 1737 *km*^2^ (or the initiation box) which increases total coral cover on the GBR by 175 *km*^2^ compared with no control after 100 years. The least effective strategy, controlling every reef on the GBR with 12.2% effort is only able to increase total coral cover on the GBR by 72 *km*^2^ after 100 years. Thus, the most effective control scenario can protect approximately 2.4 times more coral cover across the GBR than the least effective scenario after 100 years. Coral cover within the initiation box mirrors these trends, where the most effective scenario at increasing total coral cover is targeting the initiation box with 100% effort (Figure 3c).

### 3.3 Spatial spread and intensity of coral cover increase on the GBR

Culling 100% of adult COTS within the initiation box is the most effective strategy if the goal is to increase the total coral cover across the GBR. This control strategy increases coral cover on 99% of the reefs on the GBR compared to no control, or on 1696 of the 1705 reefs, by up to 18% of the reef area (Figure 4a). However, on 1486 of these reefs, the increase in coral cover is less than 1% of the reef area. The remaining 210 reefs (or 12%) of the reefs on the GBR see an increase in coral cover of 1% or more, and are mainly located on the reefs within and surrounding the outbreak initiation box, where the control effort is located (Figure 4b). Thus, although the 100% effort on the initiation box scenario is the most effective at increasing total coral cover on the GBR, the spatial extent of coral cover increase is limited to the mid-region reefs of the GBR.

**Figure 2:**
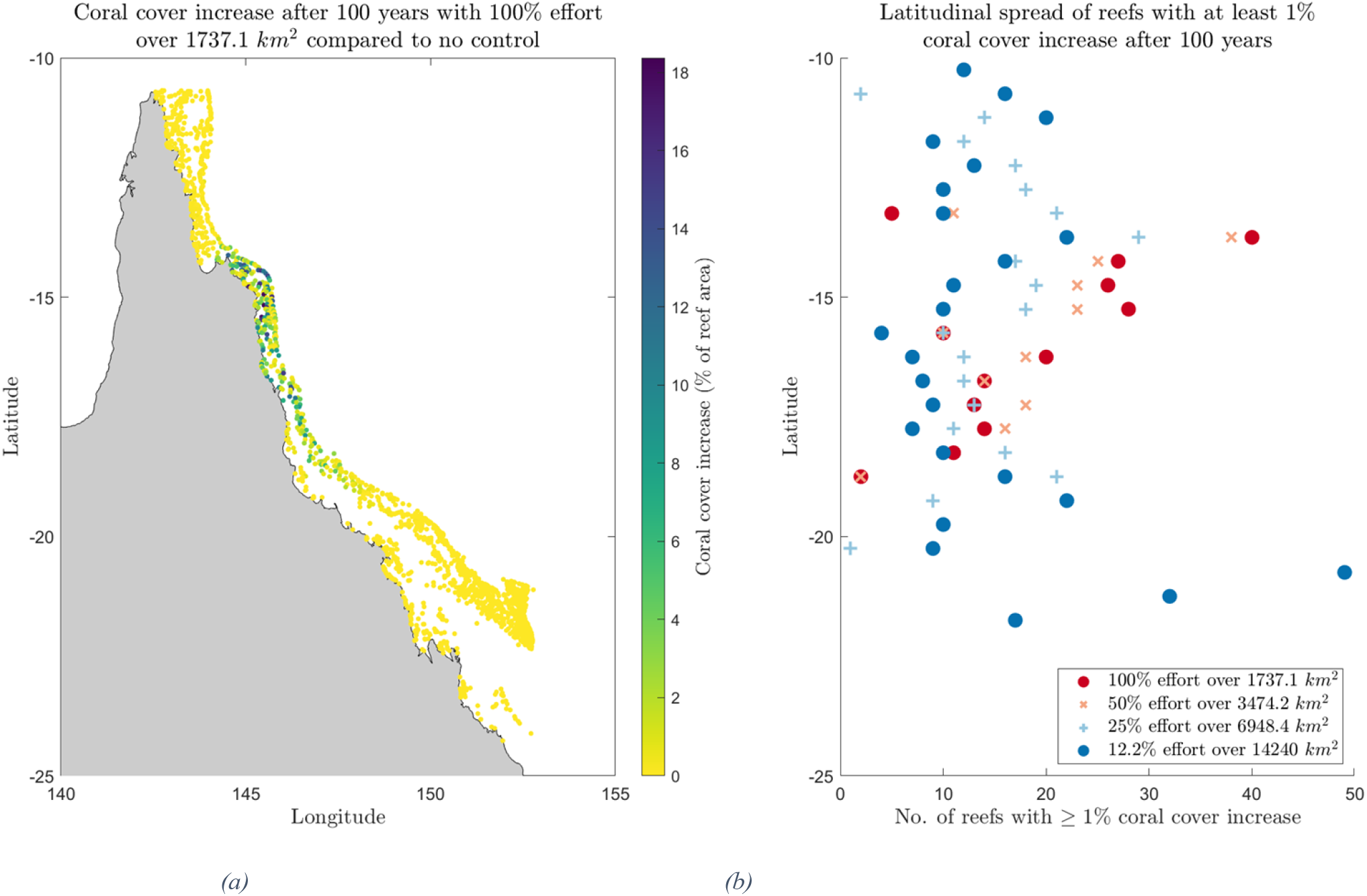
Spatial impacts on coral cover from COTS control strategies 100 years after outbreak. (a) Map of coral cover increase on the GBR for 100% effort over 1737.1 km^2^ compared to no control. Colour of each reef corresponds to the increase in coral cover, as a percentage of reef area. (b) Latitudinal spread (north-south spread) of reefs with at least 1% increase in coral cover compared to no control for control scenarios: 100% effort over 1737.1 km^2^ reefs (dark red dots), 50% effort over 3474.2 km^2^ (light red crosses), 25% effort over 6948.4 km^2^ (light blue plus signs), and 12.2% effort over 14240 km^2^ (dark blue dots).

Increasing the spatial spread of control increases the spatial extent of reefs with at least 1% increase in coral cover after 100 years compared to no control (Figure 4b). Thus, spreading the control effort wider on the GBR results in a lower increase in total coral cover compared to no control, but a higher latitudinal spread of reefs with a high increase in coral cover. If the goal is to protect coral cover across a larger spatial area on the GBR, then the most effective control strategy is to cull over the entire area of the GBR. With this control strategy, 350 reefs (or 21%) of the reefs on the GBR see an increase in coral cover of 1% or higher, but the largest increase in coral cover at a single reef is only 4%.

The choice of control strategy varies when considering different optimisation objectives. When minimising the number of reefs with 10% coral cover or less – reefs classified as ‘low coral cover’ by the Australian Institute of Marine Sciences (2021) – both 25% effort over 6948 *km*^2^, and 12.2% effort over 14240 *km*^2^ (or the entire GBR) are top performers (minimising the value of the yellow bar, Figure 5) as there are 5 fewer reefs in this category compared to no control. Alternatively, if we were to apply the maximin strategy and maximise the number of reefs with greater than 75% coral cover (maximising the value of the purple bar, Figure 5), the most effective control strategy would be 100% effort over 1737 *km*^2^ as there are 13 additional reefs within this category compared to no control.

**Figure 3:**
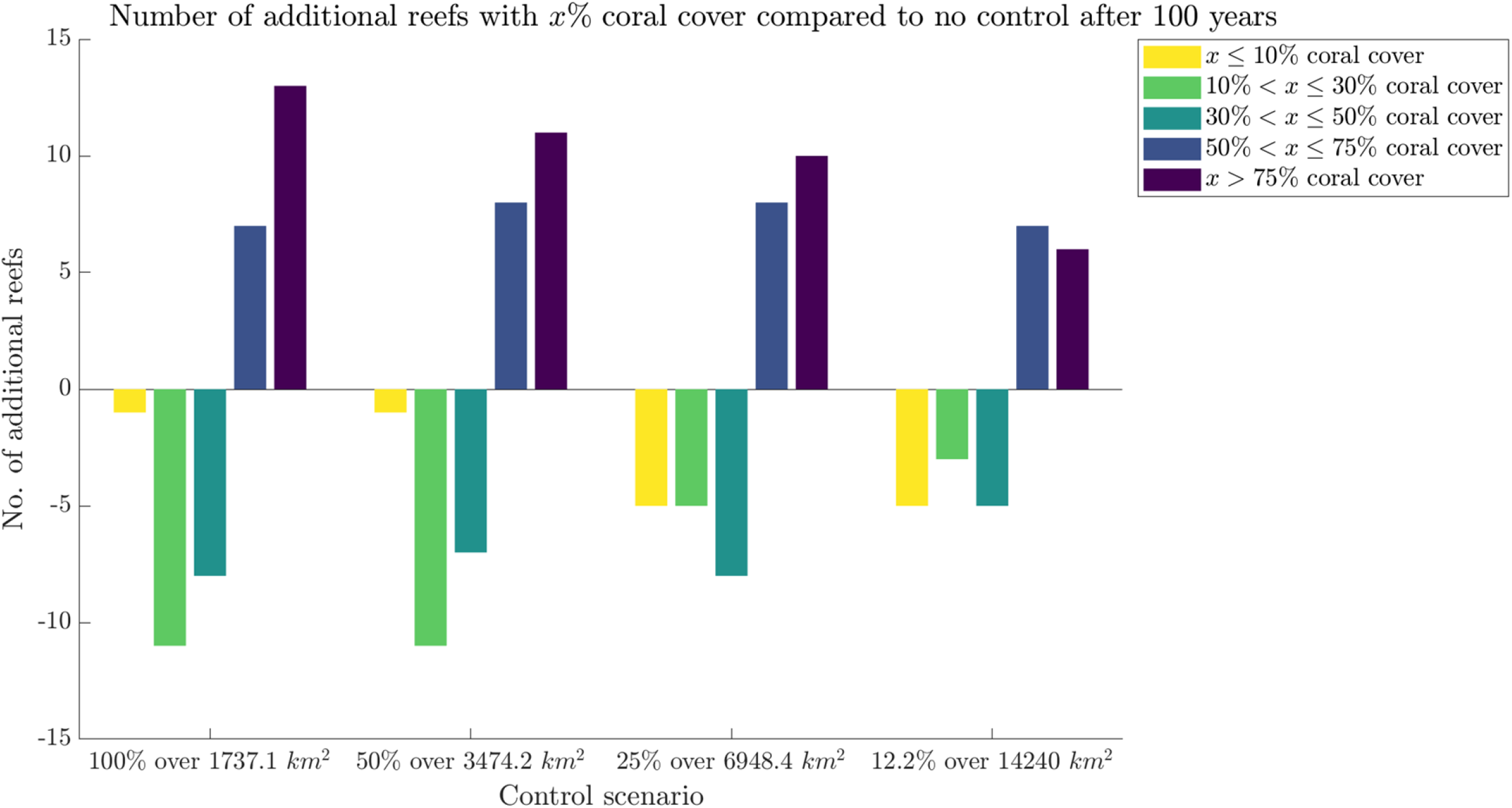
Number of reefs with different coral covers compared to no control scenario. The number of additional reefs with x% coral cover for the various control scenarios compared to no control effort, where the colour of the bar corresponds to the reef’s coral cover, as a percentage of reef area: reefs with x > 75% coral cover (dark purple), reefs with 50% < x ≤ 75% coral cover (dark blue), reefs with 30% < x ≤ 50% coral cover (light blue), reefs with 10% < x ≤ 30% coral cover (green), and reefs with x ≤ 10% coral cover (yellow). The x-axis shows the four control scenarios: 100% effort over 1737.1 km^2^ reefs, 50% effort over 3474.2 km^2^, 25% effort over 6948.4 km^2^, and 12.2% effort over 14240 km^2^ (or the entire GBR).

## 4 Discussion

To investigate the impact of various COTS control strategies on GBR coral coverage, we developed a predator-prey metacommunity model based on Morello et al. (2014), and simulated a COTS outbreak and a control response. We found that the most effective strategy was to target the source of the COTS outbreak i.e., to cull as many adult COTS as possible within the initiation box. This strategy was the most effective at increasing total coral cover (compared to no control), with an increase of 175 *km*^2^ of live coral after 100 years. Regardless of the outbreak size, targeting the source of the COTS outbreak will be beneficial for total coral cover on the GBR.

We also found that increasing the spatial extent of control effort delivered benefits to a wider spatial area, but a lower total increase compared to no control. There is therefore a trade-off between increasing the total coral cover, and expanding the range of reefs that benefit. So, if the goal is to protect a larger spatial area of coral cover across the GBR, then spreading the control effort is more effective, but if the goal is to increase total coral cover on the GBR, then targeting the source of the outbreak is more effective.

Current literature suggests that outbreaks of COTS on the GBR begin in the initiation box and thereafter spread south (Reichelt et al., 1990; Vanhatalo et al., 2017). By contrast, our simulated outbreaks spread both north and south. In addition to the expected high populations of adult COTS on southern reefs, large outbreaks were also observed in the northern GBR. Our connectivity model therefore implies that the spatial spread of COTS outbreaks moves both north and south (Reichelt et al., 1990; Vanhatalo et al., 2017).

Our model is not a predictive model, and the aim is not predict the absolute size of COTS or coral populations into the future for the given control strategies. In addition, although we identified the best strategy from our four control scenarios, this is not necessarily the optimal control strategy for the GBR. For example, we have not considered interacting factors that can increase COTS impacts such as heterogenous fishing pressures, nearby urbanization (Milne *et al*., 2023), or water quality (Matthews *et al*., 2020). Rather, we have attempted to explore how spreading the spatial extent of COTS control can affect coral and COTS populations on the GBR and found that this can have a significant impact on the populations, particularly on the spatial spread of populations.

As with any large-scale ecosystem models, we needed to make a range of assumptions when parameterising this model. Firstly, the control scenarios simulated using our model are simplistic. The cost of culling COTS does not increase linearly with the number of COTS culled; the less COTS there are at a reef, the more difficult it is to find and cull COTS. The cost of culling large proportions of COTS populations will vary from reef to reef, depending on the population size and habitat at that reef. There are also many costs associated with sending human divers to reefs, and thus the location of each reef will also have an impact on the cost; reefs that are harder to get to will be more expensive to cull at. It is also infeasible to be culling at the number of reefs we have simulated; there is inadequate funding for this (Westcott et al., 2016). However, by simulating the extremes of culling, our results range across the scope of potential actions. Furthermore, in our model we only incorporate culling of adult COTS, but in reality culling will target multiple size classes. We assume that adult culling occurs after they reproduce. This has a major impact on our results: even when we culled 100% of adults, all were allowed to reproduce first, leading to higher average populations. In reality, when culling sometimes the adult COTS will have reproduced already and other times not (Dumas et al., 2016).

The spatial distribution of COTS control can strongly impact the amount and extent of coral cover on the GBR, and is therefore an important factor when choosing control strategies. Any efficiency gains in COTS control will prove to be beneficial to the coral health and biodiversity of the GBR, particularly as pressures from climate change accelerate throughout the century.

## References

Australian Institute of Marine Sciences (2021). Reef in recovery window after decade of disturbances. Annual Summary Report of Coral Reef Condition 2020/2021.

Babcock, R. C., Milton, D. A., & Pratchett, M. S. (2016). Relationships between size and reproductive output in the crown-of-thorns starfish. Marine Biology, 163(11), 1–7.

Babcock, R. C., Plagányi, É. E., Condie, S. A., Westcott, D. A., Fletcher, C. S., Bonin, M. C., & Cameron, D. (2020). Suppressing the next crown-of-thorns outbreak on the Great Barrier Reef. Coral Reefs, 39(5), 1233–1244.

Birkeland, C. and Lucas, J. (1990). Acanthaster planci: major management problem of coral reefs. CRC press.

Bode, M., Armsworth, P. R., Fox, H. E., & Bode, L. (2012). Surrogates for reef fish connectivity when designing marine protected area networks. Marine Ecology Progress Series, 466, 155–166.

Castro-Sanguino, C., Bozec, Y. M., Condie, S. A., Fletcher, C. S., Hock, K., Roelfsema, C., … & Mumby, P. J. (2023). Control efforts of crown-of-thorns starfish outbreaks to limit future coral decline across the Great Barrier Reef. Ecosphere, 14(6), e4580.

Chen, C.-M., Drovandi, C. C., Keith, J. M., Anthony, K., Caley, M. J., and Mengersen, K. (2017). Bayesian semi-individual based model with approximate bayesian computation for parameters calibration: Modelling crown-of-thorns populations on the great barrier reef. Ecological Modelling, 364:113– 123.

Condie, S., Hepburn, M., and Mansbridge, J. (2012). Modelling and visualisation of connectivity on the great barrier reef. Proceedings of the 12th International Coral Reef Symposium, pages 9–13.

Condie, S. A., Plagányi, E. E., Morello, E. B., Hock, K., and Beeden, R. (2018). Great barrier reef’ recovery through multiple interventions. Conservation Biology, 32:1356–1367.

De’ath, G., Fabricius, K. E., Sweatman, H., and Puotinen, M. (2012). The 27-year decline of coral cover on the great barrier reef and its causes. Proceedings of the National Academy of Sciences, 109:17995– 17999.

Dietzel, A., Bode, M., Connolly, S. R., and Hughes, T. P. (2020). Long-term shifts in the colony size structure of coral populations along the great barrier reef. Proceedings of the Royal Society B, 287:20201432.

Dumas, P., Moutardier, G., Ham, J., Kaku, R., Gereva, S., Lefèvre, J., and Adjeroud, M. (2016). Timing within the reproduction cycle modulates the efficiency of village-based crown-of-thorns starfish removal. Biological Conservation, 204:237–246.

Fisher, R., O’Leary, R. A., Low-Choy, S., Mengersen, K., Knowlton, N., Brainard, R. E., and Caley, M. J. (2015). Species richness on coral reefs and the pursuit of convergent global estimates. Current Biology, 25:500–505.

Fletcher, C., Bonin, M., and Westcott, D. (2020). An ecologically-based operational strategy for COTS control: integrated decision making from the site to the regional scale. Report to the National Environmental Science Programme.

Fletcher, C. S., Castro-Sanguino, C., Condie, S., Bozec, Y. M., Hock, K., Gladish, D. W., … & Westcott, D. A. (2021). Regional-Scale Modelling Capability for Assessing Crown-of-Thorns Starfish Control. Strategies on the Great Barrier Reef.

Great Barrier Reef Marine Park Authority 1998. Great Barrier Reef Features (Version 1.4) [Dataset] 2164DB88-FD79-449E-920F-61C37ADE634B. Retrieved from http://www.gbrmpa.gov.au/geoportal.

Heenaye-Mamode Khan, M., Makoonlall, A., Nazurally, N., & Mungloo-Dilmohamud, Z. (2023). Identification of crown of thorns starfish (COTS) using convolutional neural network (CNN) and attention model. PLoS One, 18(4), e0283121.

Hock, K., Wolff, N. H., Condie, S. A., Anthony, K. R., & Mumby, P. J. (2014). Connectivity networks reveal the risks of crown-of-thorns starfish outbreaks on the Great Barrier Reef. Journal of applied ecology, 51(5), 1188–1196.

Hock, K., Wolff, N. H., Ortiz, J. C., Condie, S. A., Anthony, K. R., Blackwell, P. G., and Mumby, P. J. (2017). Connectivity and systemic resilience of the great barrier reef. PLoS biology, 15:e2003355.

Hughes, T. P., Baird, A. H., Bellwood, D. R., Card, M., Connolly, S. R., Folke, C., Grosberg, R., Hoegh-Guldberg, O., Jackson, J. B., Kleypas, J., et al. (2003). Climate change, human impacts, and the resilience of coral reefs. Science, 301:929–933.

Hughes, T. P., Kerry, J. T., Álvarez-Noriega, M., Álvarez-Romero, J. G., Anderson, K. D., Baird, A. H., Babcock, R. C., Beger, M., Bellwood, D. R., Berkelmans, R., et al. (2017). Global warming and recurrent mass bleaching of corals. Nature, 543:373–377.

Knowlton, N., Brainard, R. E., Fisher, R., Moews, M., Plaisance, L., and Caley, M. J. (2010). Coral reef biodiversity. Life in the world’s oceans: diversity distribution and abundance, pages 65–74.

Lucas, J. S. (1984). Growth, maturation and effects of diet in Acanthasterplanci (L.)(Asteroidea) and hybrids reared in the laboratory. Journal of Experimental Marine Biology and Ecology, 79(2), 129–147.

Matthews, S. A., Mellin, C., & Pratchett, M. S. (2020). Larval connectivity and water quality explain spatial distribution of crown-of-thorns starfish outbreaks across the Great Barrier Reef. In Advances in Marine Biology (Vol. 87, No. 1, pp. 223–258). Academic Press.

Matthews, S.A. (2024) Protecting Great Barrier Reef resilience through effective management of Crown-ofThorns Starfish outbreaks. PLoS One, in press

Milne, R., Anand, M., & Bauch, C. T. (2023). Preparing for and managing crown-of-thorns starfish outbreaks on reefs under threat from interacting anthropogenic stressors. Ecological Modelling, 484, 110443.

Morello, E. B., Plagányi, É. E., Babcock, R. C., Sweatman, H., Hillary, R., & Punt, A. E. (2014). Model to manage and reduce crown-of-thorns starfish outbreaks. Marine Ecology Progress Series, 512, 167–183.

Osborne, K., Dolman, A. M., Burgess, S. C., and Johns, K. A. (2011). Disturbance and the dynamics of coral cover on the great barrier reef (1995–2009). PloS One, 6:e17516.

Osborne, K., Thompson, A. A., Cheal, A. J., Emslie, M. J., Johns, K. A., Jonker, M. J., Logan, M., Miller, I. R., and Sweatman, H. P. (2017). Delayed coral recovery in a warming ocean. Global Change Biology, 23:3869–3881

Peterson, M. (2017). An Introduction to Decision Theory. Cambridge University Press.

Plagányi, É. E., Ellis, N., Blamey, L. K., Morello, E. B., Norman-Lopez, A., Robinson, W., … & Sweatman, H. (2014). Ecosystem modelling provides clues to understanding ecological tipping points. Marine Ecology Progress Series, 512, 99–113.

Plagányi, E. E., Babcock, R. C., Rogers, J., Bonin, M., and Morello, E. B. (2020). Ecological analyses to inform management targets for the culling of crown-of-thorns starfish to prevent coral decline. Coral Reefs, 39:1483–1499.

Reichelt, R., Bradbury, R., and Moran, P. (1990). Distribution of acanthaster planci outbreaks on the great barrier reef between 1966 and 1989. Coral Reefs, 9:97–103.

Vanhatalo, J., Hosack, G. R., and Sweatman, H. (2017). Spatiotemporal modelling of crown-of-thorns starfish outbreaks on the great barrier reef to inform control strategies. Journal of Applied Ecology, 54:188–197.

Westcott, D. and Fletcher, C. (2018). How effective are management responses in controlling crown of-thorns starfish and their impacts on the great barrier reef. Report for NESP Tropical Water Quality Hub, Cairns.

Westcott, D., Fletcher, C., Babcock, R., and Plagányi-Lloyd, E. (2016). A strategy to link research and management of crown-of-thorns starfish on the great barrier reef: An integrated pest management approach. Report to the National Environmental Science Programme, 77.

